# Characterizing particle dynamics in live imaging through stochastic physical models and machine learning

**DOI:** 10.1101/2024.12.17.628916

**Authors:** G. Nardi, M. Santos Sano, M. Bilay, A. Brelot, J.-C. Olivo-Marin, T. Lagache

## Abstract

Particle dynamics determine the orchestration of molecular signaling in cellular processes. A wide range of subdiffusive motions has been described at the cell interior and membrane, corresponding to different environmental constraints. However, the standard methods for motion analysis, embedded in a diffusion-based framework, lack robustness for capturing the complexity of stochastic dynamics. This work develops a classification method to detect the five main stochastic laws modeling particle dynamics accurately. The method builds on machine-learning techniques that use features properly designed to capture the intrinsic geometric properties of trajectories governed by the different processes. This guarantees the accurate classification of observed dynamics in an interpretable and explainable framework. The main asset of this approach is its capability to distinguish different subdiffusive behaviors making it a privileged tool for biological investigations. The robustness to localization error and motion composition is proven, ensuring its reliability on experimental data. Moreover, the classification of composed trajectories is investigated, showing that the method can uncover the path’s mono-vs bi-dynamics nature. The method is used to study the dynamics of membrane receptors CCR5, involved in HIV infection. Comparing the basal state to an agonist-bound state which displays potent anti-HIV-1 activity, we show that the latter affects the natural dynamic state of receptors, thus clarifying the link between movement and receptor activation.

## 1 Introduction

Cell processes are governed by inter-and extra-cellular signaling triggered by the dynamic interaction of different molecules. Live cell imaging with fluorescence microscopy enables the analysis of the dynamics of targeted particles (e.g., molecules, receptors, and viruses) facilitating the study of biological processes at a sub-cellular scale. In this context, particle tracking and related motion analysis are of fundamental importance in revealing the impact of the cell environment and the interaction with other molecules. This enables the observation of the stochastic trajectories of viruses inside the cell cytoplasm, highlighting the important role of the cytoskeleton for virus transport [1, 2, 3]. Similarly, live imaging of cell receptors helped to understand the activation pathways [4], the organization of the membrane in functional microdomains [5] and endosomal routing [6].

Although several methods [7, 8] enable the trajectory reconstruction of labeled molecules in time-lapse fluorescence imaging, motion classification lacks robust methods for capturing the complexity of stochastic dynamics. Motion classification is usually developed within a diffusion-based framework enabling the detection of three classes of motion [9, 4, 10]: *Brownian* motion modeling free diffusion of objects [11], *subdiffusive* motion describing movements constrained by local crowding and physical interactions within cell microdomains [12, 13], and *superdiffusive* motion that accounts for active transport such as molecular cargos along the cell cytoskeleton [2, 14, 15]. However, main cellular mechanisms are governed by co-existing *subdiffusive* behaviors [16] generating particle trapping in a potential field [17], constrained evolution due to the viscoelastic properties of the cellular medium [18], or confinement within nanodomains [19]. Therefore, the inference of the stochastic laws governing these different behaviors [20, 21, 22] represents an important challenge for motion classification allowing a finer description of particle dynamics which is not feasible within the diffusion-based framework.

The second challenge concerns the definition of robust descriptors discriminating the different motion classes. The diffusion-based classification traditionally uses the mean squared displacement (MSD) function, discriminating the diffusion class based on its polynomial order [23, 24]. However, the MSD criterion has many drawbacks, linked to the accurate statistical estimate of the fitting coefficient (see Appendix A), which has encouraged the introduction of other analytical descriptors [25, 26]. Such descriptors include features related to the Gaussianity of displacement distribution [27] and specific trajectory features, such as the standardized longest distance traveled by a particle [9,4, 28], or the *p*-variation for a refined description of some *subdiffusive* processes [20]. Classification algorithms build on these features to define statistical hypothesis tests [9, 20, 4, 28] or machine learning (ML) approaches [29, 30] in order to perform diffusion-based classification [30] or distinguishing specific dynamics [20, 29, 10]. To avoid the definition of *ad hoc* features, deep-learning approaches have recently been developed [31, 22], achieving higher accuracy at the cost of standard interpretability issues that limit understanding of studied phenomena.

The present work deals with the previous challenges, introducing a novel model to characterize the wide variety of cellular dynamics based on explainable well-suited features. The proposed model classifies the stochastic laws governing the trajectory and can differentiate, in addition to standard Brownian (BM) and directed motion (DIR), three additional subdiffusive and non-ergodic dynamics: 1) the Ornstein–Uhlenbeck (OU) process that models the attraction of diffusing objects towards a potential well [32], 2) the fractional Brownian motion (FBM) that describes objects’ motion in a crowded and constrained environment like the viscous cellular medium [33], and 3) the Continuous-Time Random Walk (CTRW) that accounts for non-ergodic evolutions alternating jumps and pauses due to trapping by molecular obstacles [34]. Several analytical descriptors have been proposed in the literature to characterize these different processes [20, 29, 10], but none of the existing methods can distinguish such a wide family of motions. To overcome these limitations, the present work proposes well-suited geometrical features of trajectories that describe how particles unfurl in space and encode the intrinsic properties of each process. However, instead of defining analytical criteria or statistical hypothesis tests, our method builds on the versatile Random Forest framework allowing an accurate classification of the previous five processes and proving the power of the machinelearning approach for motion analysis. Moreover, our feature-based approach guarantees the interpretability of tracked objects’ complex dynamics, facilitating the inference of the underlying biophysics.

After a thorough validation on synthetic data, we used our method to analyze a set of C-C chemokine receptor type 5 (CCR5) trajectories at the cell membrane obtained with total internal reflection fluorescence (TIRF) microscopy. We point out that TIRF enables observation at 200*nm* from the coverslip allowing access to events that happen at the plasma membrane. CCR5 is critically important in the context of human immunodeficiency virus (HIV) infection serving as a virus’ co-receptor to enter and infect host cells.

We analyzed the CCR5 trajectories using our features-based machine-learning method revealing a complex landscape characterized by subdiffusive dynamics. At basal state, the majority of tracks exhibit either intermittent motion (CTRW) or continuous dynamics within a constrained environment (FBM); on the other hand, after stimulation with PSC-RANTES, an analog of the natural ligand RANTES/CCL5 which displays potent anti-HIV-1 activity, we observed a decrease of CTRW dynamics in favor of FBM motion. Thanks to our method, we were able to reveal the impact of PSC-RANTES stimulation on receptors’ behavior, prompting them to move from free movement, characterized by jumps and pauses, to constrained motion at the cell membrane aiming for internalization. This highlights the importance of distinguishing a larger set of stochastic dynamics, especially in the subdiffusive regime, to elucidate the link between movement and receptor activation in the orchestration of cell signaling.

## 2 Methods

In this section, we present the main steps involved in setting the proposed method. First, Section 2.1 briefly recalls how discrete stochastic trajectories were simulated to define a dataset used to train and validate the proposed method. Section 2.2 introduces the geometric features used in this work to characterize the different motion classes. Finally, Section 2.3 defines the supervised approach used for classification.

## 2.1 Simulating discrete stochastic dynamics

We consider the five stochastic processes modeling the main particle behaviors observed in cellular dynamics (see Appendix A for details): 1) free motion characterized by Brownian trajectories (BM); 2) confined and local movements generated by the Ornstein-Uhlhenbeck process (OU); 3) directed and superdiffusive motion governed by a drift (DIR); 4) constrained evolutions due to a crowded and viscous cellular medium described by the fractional Brownian Motion model (FBM); 5) movements alternating jumps and pauses describing trapping phenomena and modeled by the Continuous-Time Random Walks (CTRW).

These five stochastic laws can be described theoretically using stochastic differential equations. The discretization of the related equations enables thus the simulation of discrete trajectories used for machine-learning method training and validation. Appendix A gives a complete presentation of the mathematical properties of the considered processes and describes how their numerical simulation is performed to generate synthetic datasets of stochastic trajectories. These datasets are used in the rest of the paper for illustrating the features introduced in Section 2.2 and for method training and validation in Sections 2.3 and 3.1.

Simulated trajectories consistently exhibit the dynamic behavior induced by the related process (e.g., confinement, trapping, constrained evolutions). This results in trajectories with characteristic spatial evolutions whose geometric properties allow inferring the corresponding process. This justifies the features-based approach used in this work, which is based on geometrical descriptors of trajectories encoding the different dynamic characters and enabling process discrimination.

### 2.2 Designing trajectories’ geometrical features

In this work, we consider two families of features, the first accounting for directionality and the second quantifying the spreading character of particles.

Directionality can be described using the distribution of angles between successive positions, encoding the persistent, Brownian, or antipersistent character of trajectories [27]. This translates into our framework identifying paths driven by restoring forces (majority of large angles, corresponding to OU or FBM (*H <* 1*/*2)), propagation behavior (majority of small angles, corresponding to DIR or FBM (*H >* 1*/*2)) and random motion (BM is characterized by uniform-like angle distribution). On the other hand, due to trapping, most of the angles between successive positions are null in CTRW paths.

The second set of features describes the spreading properties of particles based on Ripley’s indices on concentric balls (see equation (2.3)). This allows describing how the particle unfurls in space and helps, in particular, to depict confined ( OU) a nd spreading (DIR) motions. To our knowledge, this work is the first to use t his family of features for motion

In the following, we denote a discrete trajectory as a set of *N* positions *X*_*i*_ ∈ ℝ^2^, with independent components, corresponding to different times *t*_1_ *<* … *< t*_*N*_ ∈ ℝ^+^:

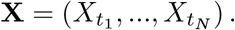

#### Description of trajectory directionality

For every three consecutive points of the trajectory 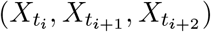, we consider the angle [27]:

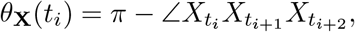

which is considered as positive if 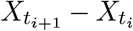 rotates on 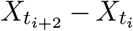 in a counterclockwise way. In the case of CTRW, most angles are null since the particle often remains stationary at the same point. Therefore, non-null angles are computed between triplets of not necessarily consecutive displacements.

As shown in Figure 1, the density 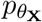 of angle distribution exhibits a different analytical dependence for different processes. Brownian motion has a uniform distribution of angles because of its random displacements [27]. On the other hand, a motion governed by a drift exhibits displacements around the privileged direction resulting in small angles between successive positions; then, the histogram of the angles can be approximated by a Gaussian distribution centered at zero. Finally, in confined motions driven by restoring forces, most of the angles must be larger than *π/*2 allowing the particle to go back to the equilibrium point. This suggests that the convexity of the angle distribution gives a first approximation of the histogram shape allowing for distinguishing different dynamics.

**Figure 1.**
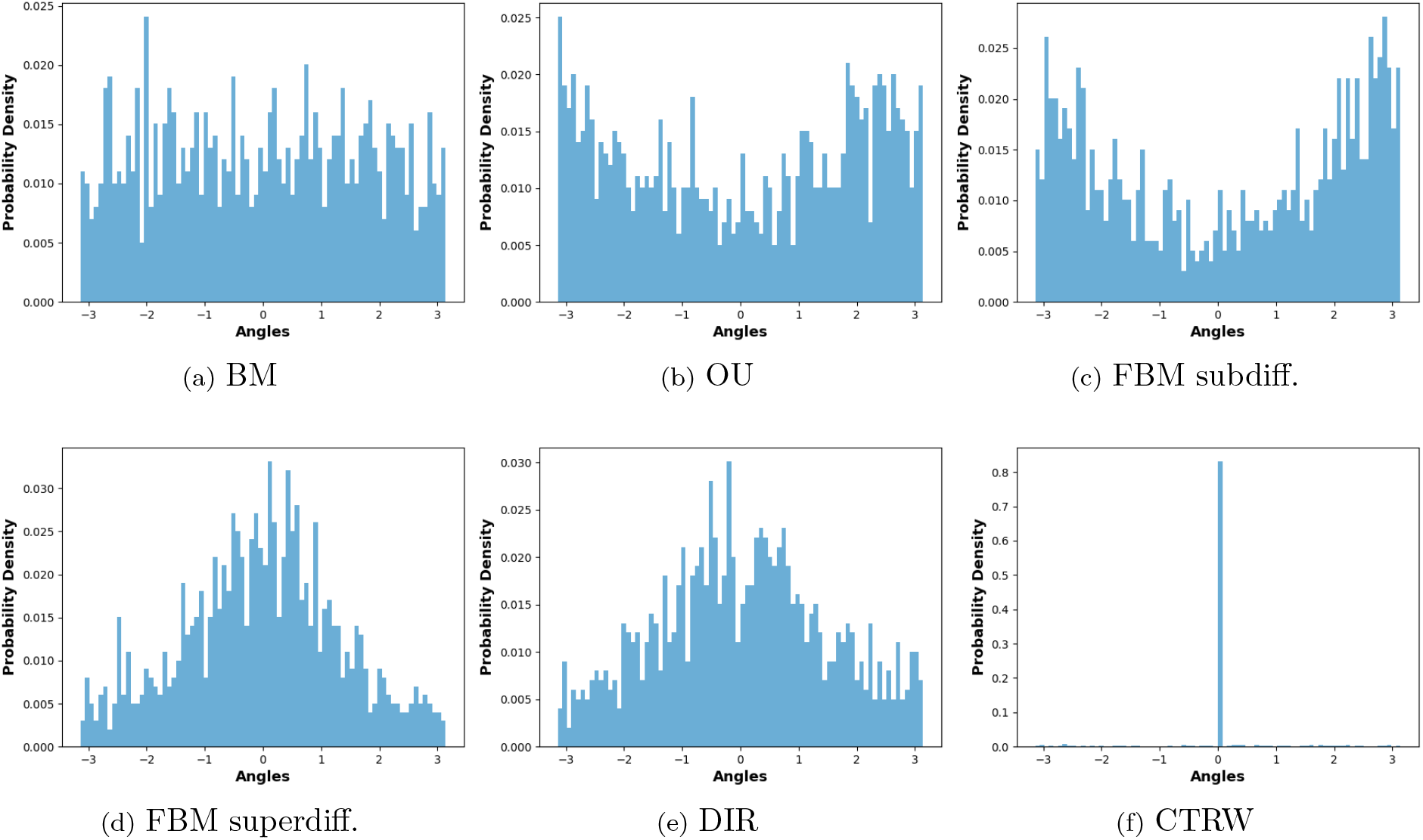
Examples of angle distributions for different processes: (a) BM (*σ* = 1); (b) OU (*λ* = 0.5, *σ* = 1); (c) FBM with *H* = 0.2; (d) FBM with *H* = 0.8; (e) DIR ( ∥*u*∥ = 0.7, *σ* = 1); (f) CTRW with *σ* = 1 and *γ* = 0.9.

Thus, to estimate the convexity of the distribution, the following fitting is performed:

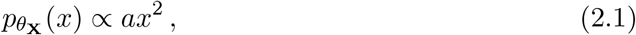

and the value of *a* defines a geometrical feature describing the angle variability.

Moreover, to establish the existence of a preferred motion direction along the trajectory we also consider the following index of directionality [27]:

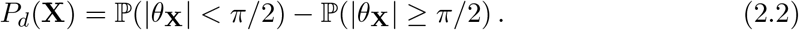

A positive *P*_*d*_ indicates that most of the angles are smaller than *π/*2 resulting in a persistent movement along a privileged direction (corresponding to diffusive dynamics). In contrast, a negative *P*_*d*_ indicates an anti-persistent movement exhibiting larger angles (corresponding to a confined dynamic governed by restoring forces).

#### Encoding trajectory spreading

The spreading character of a trajectory can be described by analyzing its displacements through concentric balls. This enables capturing (local) confinement or constant drift and quantifying the motion amplitude.

To this goal, we consider the Ripley’s index *K*_*r*_ in a ball *B* 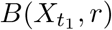 of radius *r* centered at the starting point [35]:

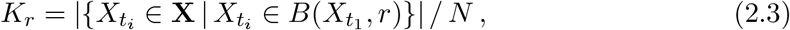

accounting for the number of trajectory points living in that ball.

To make this computation consistent with the trajectory dynamics, we define the reference radius:

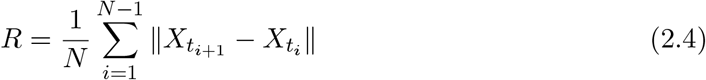

and we compute the vector:

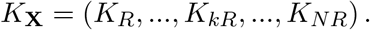

Figure 2 shows several vectors *K*_**X**_ for different processes suggesting that its analytical shape enables differentiating d ifferent dynamics.

**Figure 2.**
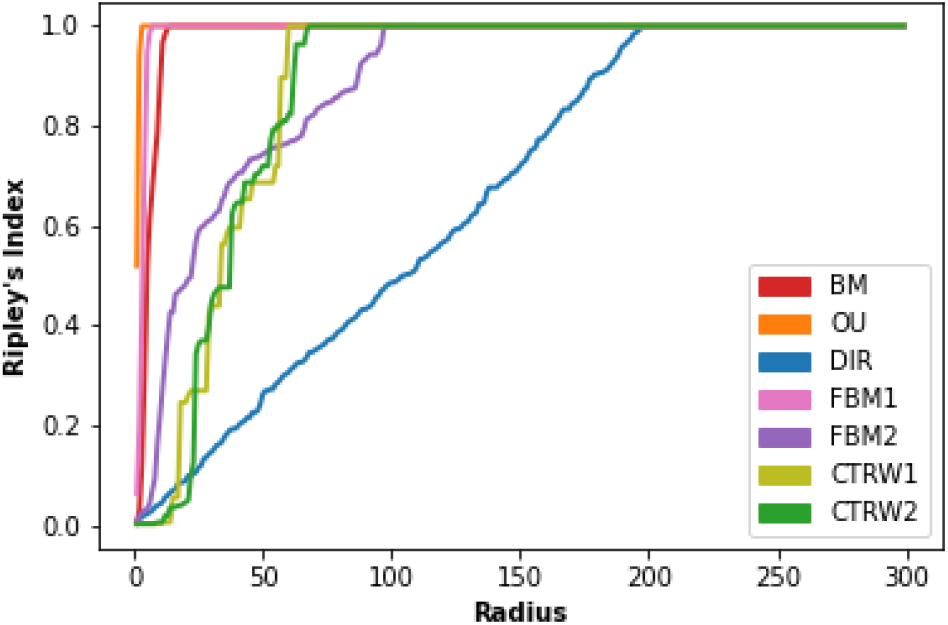
Curves of Ripley’s indices exemplified by a graph for each process (radius is expressed in terms of indices *k*s’): BM (*σ* = 1), OU (*λ* = 0.5, *σ* = 1), DIR (∥ *u* ∥ = 0.7, *σ* = 1), FBM (FBM1 with *H* = 0.2, and FBM2 with *H* = 0.8), CTRW (*σ* = 1, *γ* = 0.01 for CTRW1 and *γ* = 0.9 for CTRW2).

By definition o f R ipley’s i ndex (2.3), i t f ollows that *K* _**X**_ i s a n on-decreasing function with *K*_*r*_ = 1 if *r* is larger than the maximal distance of the particle from its initial point. This suggests estimating the analytical law of *K*_**X**_, as a function of index *k*, using the following fitting:

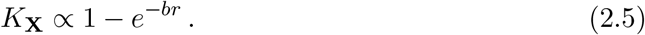

We then consider *b* as the third feature of interest. Fig. 3a displays the value of this feature for a set of trajectories simulating the different processes (see Section 2.1), highlighting how its value accurately depicts different spreading trends (e.g., confined, trapping, superdiffusive).

**Figure 3.**
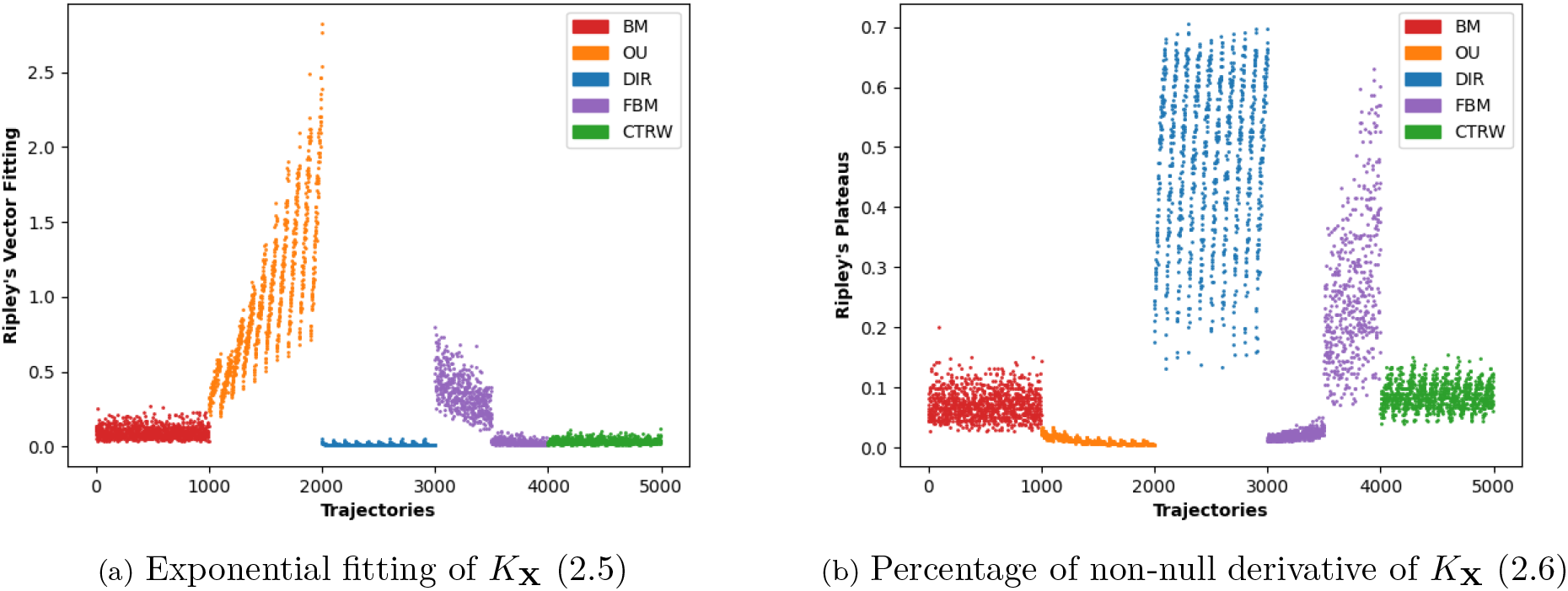
Value of features defined in equation (2.5) (a) and equation (2.6) (b) for different trajectories corresponding to the considered processes (see definition of the training set in Section 2.1).

Finally, we take into account the non-decreasing behavior of *K*_**X**_ by considering the percentage of its points with a non-null derivative:

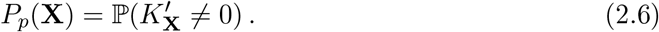

*P*_*p*_(**X**) will define the last feature considered in this work and is displayed in Fig. 3b for simulated trajectories (see Section 2.1). The null derivative for *K*_**X**_ results in local plateaus in its graph, revealing a (locally) confined movement leading to constant Ripley’s index on successive concentric balls. On the other hand, a high percentage of non-null derivatives corresponds to spreading motions, characterized by the scarcity of backward movement and driven by a privileged direction, as in the case of superdiffusive trajectories.

### 2.3 Classification method

Machine-learning methods face increasing competition from deep-learning techniques, which give accurate classification with limited preprocessing tasks like the design of *ad hoc* motion descriptors [22]. However, compared to deep-learning models, the features-based approach offers explainability and interpretability which is of fundamental importance to establish a correlation between dynamics and associated biophysical constraints. For these reasons, we chose a supervised features-based approach for this work.

The proposed method exploits the Random Forest algorithm to establish a novel approach for stochastic process discrimination. Trained on the features defined in Section 2.2 (see equations (2.1), (2.2), (2.5), and (2.6)), and computed on the entire trajectories of a given length, the algorithm accurately learns the stochastic law described by the four geometric descriptors. Training and validation are performed on independent datasets, generated as detailed in Section 2.1, using the model’s name as labels (BM, OU, DIR, FBM, CTRW).

Compared to standard classification algorithms (e.g., k-NN, SVM), the Random Forest algorithm is particularly competitive in capturing complex non-linear patterns within data. Random Forest classifiers average predictions from a set of trees (ten in our case) without any *a priori* on the data space structure. This results in a seamless approach based only on a few hyperparameters and guaranteeing interpretability and feature importance analysis.

## 3 Results

The first part of this section measures the method performances on simulated trajectories (Section 3.1). We first study the influence of localization error and trajectories’ length variability on the method’s accuracy. Then, we compare our method to standard diffusion-based classification and investigate the method’s robustness to motion composition. After having assessed the method’s accuracy and robustness for an extensive set of simulations, we apply the method to biological data and study the dynamical behavior of CCR5 receptors at the cell membrane (Section 3.2).

### 3.1 Classification of simulated trajectories

#### Method accuracy depends on trajectory length

Table 1 and Table 2 present the method results for different trajectory lengths, which proves that trajectory characteristics need time to emerge distinctively and supports the inaccuracy of shorter trajectory classification. Most misclassifications concern shorter FBM trajectories (classified as OU or BM for *N* = 40, 70) due to the variability of their dynamics behaviors depending on *H*. This error decreases significantly with increasing length, reaffirming that FBM describes intrinsically different dynamics. On the other hand, CTRW behavior is easily learned, also for short trajectories, due to successive waiting times and the properties of 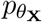 and *K*_**X**_.

**Table 1:** Results of the machine learning method: Recall by motion class is shown for different trajectory lengths.

| Length | BM | OU | DIR | FBM | CTRW |
| --- | --- | --- | --- | --- | --- |
| N=40 | 82.7 | 80 | 80 | 60.7 | 99.6 |
| N=70 | 89.9 | 89.7 | 90.2 | 80 | 100 |
| N=100 | 94.2 | 92.3 | 92.5 | 89.1 | 99.9 |
| N=200 | 98.9 | 98.2 | 97.4 | 96.3 | 100 |
| N=300 | 98.8 | 99 | 99.6 | 98.6 | 100 |

**Table 2:** Results of the machine learning method: Precision by motion class is shown for different trajectory lengths.

| Length | BM | OU | DIR | FBM | CTRW |
| --- | --- | --- | --- | --- | --- |
| N=40 | 69.9 | 81.5 | 81.7 | 70.6 | 100 |
| N=70 | 82.3 | 89.3 | 90.9 | 87.8 | 100 |
| N=100 | 88 | 92.9 | 94.1 | 93.5 | 100 |
| N=200 | 95.8 | 97.6 | 99 | 98.5 | 100 |
| N=300 | 98.4 | 98.2 | 99.5 | 99.9 | 100 |

**Table 3:** List of main properties for each stochastic process. Stationarity properties for OU process depend on the initial state, and brackets hold if 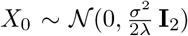; FBM refers to the case *H ≠*1/2; CTRW properties depend on the distribution of waiting times, brackets hold for exponential distribution and do not for a power-law distribution.

| Process | Continuous | Gaussian | Stationary | Stationary<br>Increments | Independent<br>Increments | Ergodic |
| --- | --- | --- | --- | --- | --- | --- |
| **BM, DIR** | × | × |  | × | × | × |
| **OU** | × | × | (×) | (×) |  | × |
| **FBM** | × | × |  | × |  | × |
| **CTRW** |  | × |  | (×) | × | (×) |

Figure 4 summarizes the method’s accuracy against the trajectory length. Through this paper, the error bars shown in graphs represent the related Binomial proportion confidence interval with confidence level set to 95%.

**Figure 4.**
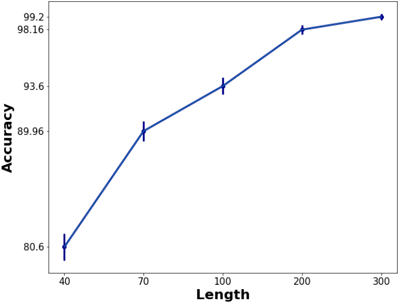
Method accuracy depending on trajectory length.

Finally, Figure 5 shows the results of studying the features’ importance for each class based on the mean decrease accuracy method. A random permutation of values is performed for each feature, and the trained model is applied to the new dataset. For each class, the difference between the original and the new recall estimates how that feature discriminates the class (because of randomness, we average the results of ten independent permutations).

**Figure 5.**
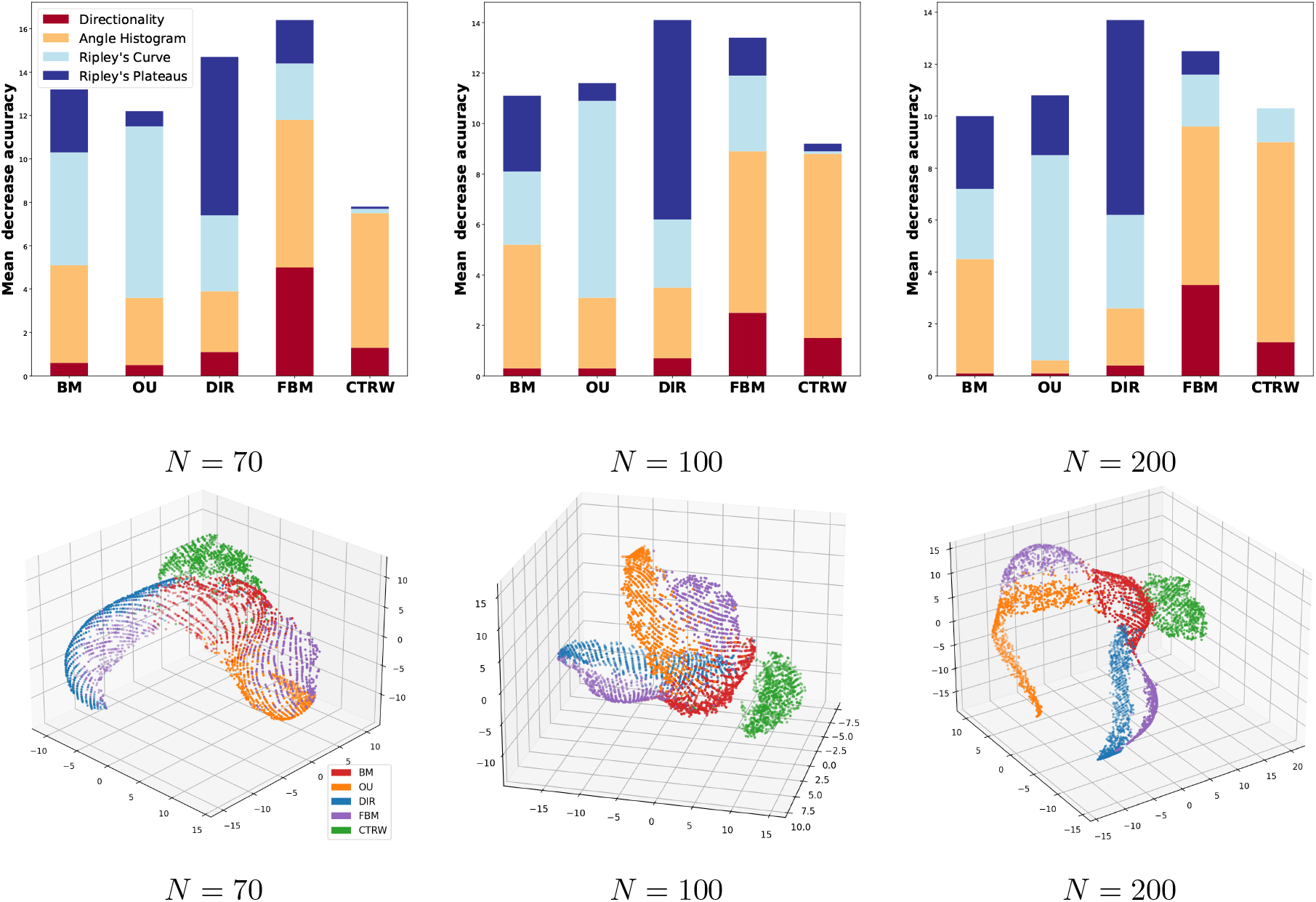
Features analysis on the train set for several trajectory lengths. Top : Feature-specific mean decrease accuracy per motion class. Bottom : tSNE representation in 3D of the feature set.

Although the importance scores vary with the trajectory length, we can point out some common trends. The fitting coefficient of Ripley’s curve is important to identify OU process; this is due to the confined behavior of OU paths resulting in a rapidly increasing Ripley’s curve. Ripley’s plateaus distinguish DIR motions whose Ripley’s curves are characterized by a non-null derivative due to the constant spreading along the trajectory driven by the drift component. CTRW is classified based on the angle histogram close to a Dirac distribution at zero for this process. A more uniform mix of features identifies BM and FBM. In particular, FBM classification is the most impacted by the directionality feature, differentiating it from its confined and directed *alter ego* (OU, DIR).

Previous results assert the dual power of the proposed approach. Firstly, the features used in this work encode the intrinsic properties of the different processes enabling the geometric characterization of several dynamic behaviors. Secondly, an accurate classification based on stochastic laws provides a finer prism for motion classification, going beyond the traditional diffusion framework and correlating with biological phenomenology.

#### Impact of localization error and immobility threshold

A critical step for tracking analysis concerns estimating the localization error (LocErr) affecting particle positions and summarizing the inaccuracies related to intensity variability in confocal videos and position computation performed by detection algorithms. The estimation of LocErr is central for trajectory analysis, in particular for methods dealing with CTRW, because it strongly affects the definition of particle immobility. However, a precise estimation of this error is a difficult task, depending on both the optical system used for fluorescence microscopy [36] and the specificities of live cell imaging, as the thermal and mechanical fluctuations of the cell membrane [37].

Then, similarly to [30, 29], to study the robustness of our method to localization error, we express LocErr as fractions of particle diffusion coefficient instead of using a fixed estimation based on physical constraints. A new dataset is defined, in which each trajectory position is modified by adding a Gaussian perturbation. We model the LocErr as a Gaussian process with zero mean and variance *σ* verifying

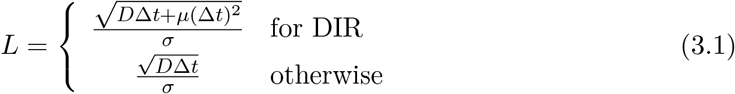

where L denotes the localization error level, *D* is the diffusion coefficient estimated via the fitting formula for the MSD ( ∝ 4*Dt*^*α*^), and *µ* is the norm of the drift component used to simulate directed paths (DIR). Ranging *L* from 1 (high error) to 9 (low error), the previous equation allows the computation of *σ* = *σ*(*L*) and the generation of datasets perturbed by different LocErr.

We tested the robustness to localization error by considering the model trained on non-perturbed trajectories of length 100 and collecting its performance on the previously perturbated datasets for different *L*.

However, in our case, this standard approach to the analysis of the impact of LocErr needs a preliminary step. Indeed, the localization error strongly impacts the immobility of particles, which is the main characteristic of CTRW paths. An immobility threshold should be applied to trajectory points to detect immobile particles over different time intervals. To do this, we compare each point with the previous one, and if their distance is smaller than the given threshold, its position is set equal to the previous one. The threshold used for this immobility correction equals 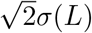, where *σ*(*L*) is the variance of the LocErr applied to the trajectory.

Figure 3.1 reports the performance results for the method tested on synthetically perturbed trajectories corrected with such a threshold. Starting at L=3, the accuracy becomes stable around the values previously obtained for the method validation.

For trajectories with high LocErr (L=1,2), CTRW and OU are well classified but the accuracy of the other motion classes is negatively affected. Comparing the recall and precision graph we can observe that BM and DIR are poorly detected (low recall) while only a few trajectories are misclassified with these motions (high precision). On the other hand, the bad recall and precision of FBM reveal that this class is poorly detected for trajectories with high LocErr and contains a high percentage of misclassification.

Finally, the low precision for CTRW points out an overestimation of immobility, for instance for BM or FBM trajectories. This is consistent with the fact that, for L=1,2, the diffusion coefficient used for non-perturbed trajectory simulation is similar to the LocErr variance and immobility threshold. For larger L, the diffusion coefficient of non-perturbed trajectories becomes larger than the LocErr variance, avoiding immobility overestimation and improving performance.

In conclusion, this analysis shows that, for LocErr ≥ 3, perturbed trajectories are classified with accuracy comparable to that one obtained in Tab.1 and 2 for *N* = 100. This proves, in particular, that the classification algorithm is robust to mean localization errors occurring in data acquisition and treatment, ensuring its reliability on experimental data.

#### Impact of length inequality

The proposed method being designed to classify trajectories with fixed length *N* its deployment faces the practical issue of collecting experimental trajectories of the same length, requiring a considerable sorting of acquired paths and, therefore, a larger number of acquisitions. Indeed, experimental videos show particles at different life stages and with independent fates, some of them leaving the field of view, others entering it. Moreover, the environment density and the fluorescence variability over time can prevent the detection of some of them over short time intervals implying a large variability in trajectory lengths.

Then it is important to study how length inequality affects the method’s performance, setting a suitable threshold on *N* for trajectory collection to facilitate the experimental acquisition tasks and the related data analysis work. To do this, we trained the proposed method on the dataset of trajectories of length 100 and tested it on several datasets of trajectories with lengths from 70 to 130. The results are reported in Figure 7.

**Figure 6.**
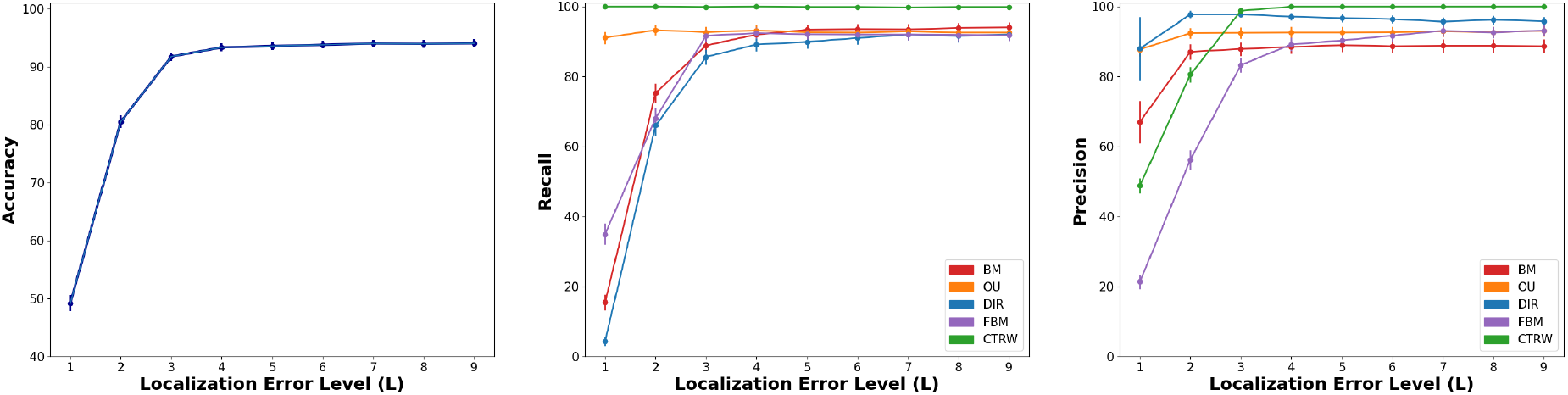
Analysis of accuracy, recall, and precision for the method trained on non-perturbed trajectories (*N* = 100, Δ*t* = 1*/*30) and tested on trajectories affected by LocErr for different localization error levels *L*. The immobility correction is performed with threshold 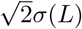. Starting at L=3, the accuracy becomes stable around the value previously obtained for the method validation. For trajectory perturbation with high LocErr (*L* = 1, 2), OU and CTRW show a high recall proving their robustness to LocErr. However, the particle immobility is overestimated resulting in poor CTRW precision. The other classes are strongly affected by high LocErr, particularly FBM, which exhibits poor performance (low recall) and high misclassification (low precision).

**Figure 7.**
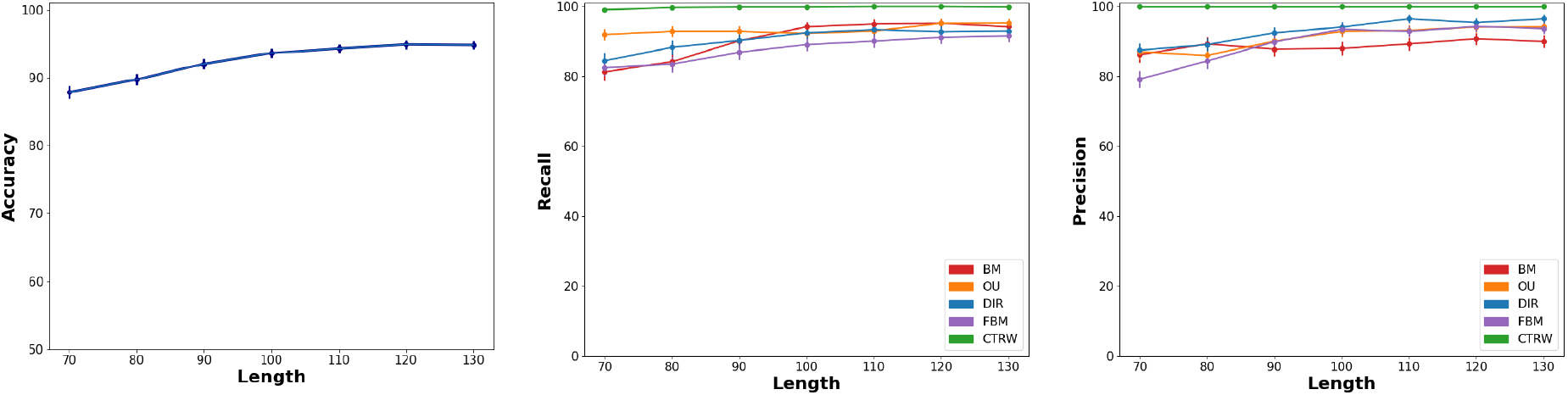
Results of accuracy, recall, and precision for the method trained on trajectories of length 100 and tested on trajectories with different lengths (Δ*t* = 1*/*30). The method’s accuracy slightly decreases whenever the trained method is applied on shorter trajectories of at most ten time points (accuracy of 93.6 % on trajectories with *N* = 100 vs accuracy of 92% on trajectories with *N* = 90). This allows for collecting reconstructed trajectories with slightly different lengths without affecting prediction accuracy.

For instance, using the model on shorter trajectories leads to misclassifying directed paths predicted as FBM ones, whereas misclassified FBM trajectories are mostly labeled as OU. Surprisingly, BM motion is the most sensitive to length variation and is mostly confused with the DIR class. OU and CTRW paths are uniformly well classified on shorter trajectories. On the other hand, results slightly improve if the model is tested on longer trajectories.

Finally, we point out that, by limiting the length variance to Δ*N* = 10 in the shortening sense (*N >* 90), we register a maximum gap of the accuracy of 1.6% compared to the accuracy on trajectories with the same length (N=100, accuracy = 93.6%, see Fig. 4). This proves the robustness of our method to slight variability of length for analyzed trajectories (up to Δ*N* = 10), allowing for collecting more data and guaranteeing the same accuracy in motion analysis.

#### Comparison to statistical hypothesis testing method

A comparison of the proposed method to the standard diffusion-based classification enables highlighting both the consistency of the two approaches and the richness of distinguishing several stochastic processes. To this goal, we consider the novel method developed in [9] performing a diffusion-based classification of paths with length *N* based on a statistical test procedure. This approach bypasses thus the statistical inaccuracy of the MSD definition which is, in particular, a critical point for the classification of short paths.

They consider the distribution of the standardized maximal distance of the particle from its starting point along its trajectory:

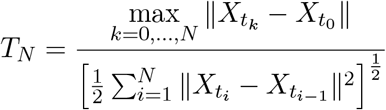

whose the quantiles *q*_2.5_ and *q*_97.5_, computed for Brownian paths (BM) of length *N*, enable setting the following three-hypothesis-test procedure:

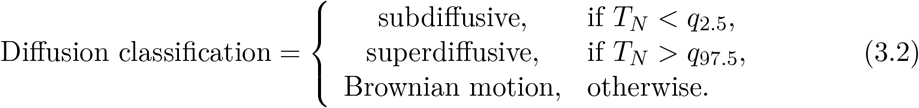

We compare the proposed method, based on process classification, to the hypothesis test (3.2), performing diffusion classification. We consider the test dataset for trajectories of length *N* = 100, and predictions are made both via our method (trained on trajectories of the same length) and via the three-decision-test (3.2) (the quantiles are computed via Monte-Carlo simulation of 100001 Brownian trajectories of length *N* = 100).

In Figure 8, we report the normalized (with respect to process classes) confusion matrices of motion classification versus motion or diffusion classes. BM, OU, and DIR are correctly classified in terms of diffusion. FBM is correctly split into two classes of subdiffusive (*H <* 1*/*2) and superdiffusive (*H >* 1*/*2) paths. However, compared to our method, a larger part of the FBM track is misclassified by the test (3.2). Misclassified paths are labeled as free motion (BM), although they have been simulated with Hurst coefficients well distinguished from 1*/*2 (*H* ∈ [0.15, 0.3] ∪ [0.7, 0.85]). Finally, the test (3.2) misclassifies the totality of CTRW trajectories as free motion. Although this is consistent with the Gaussianity of the jumps distribution, this shows that the (3.2) is unadapted to the detection of CTRW paths.

**Figure 8.**
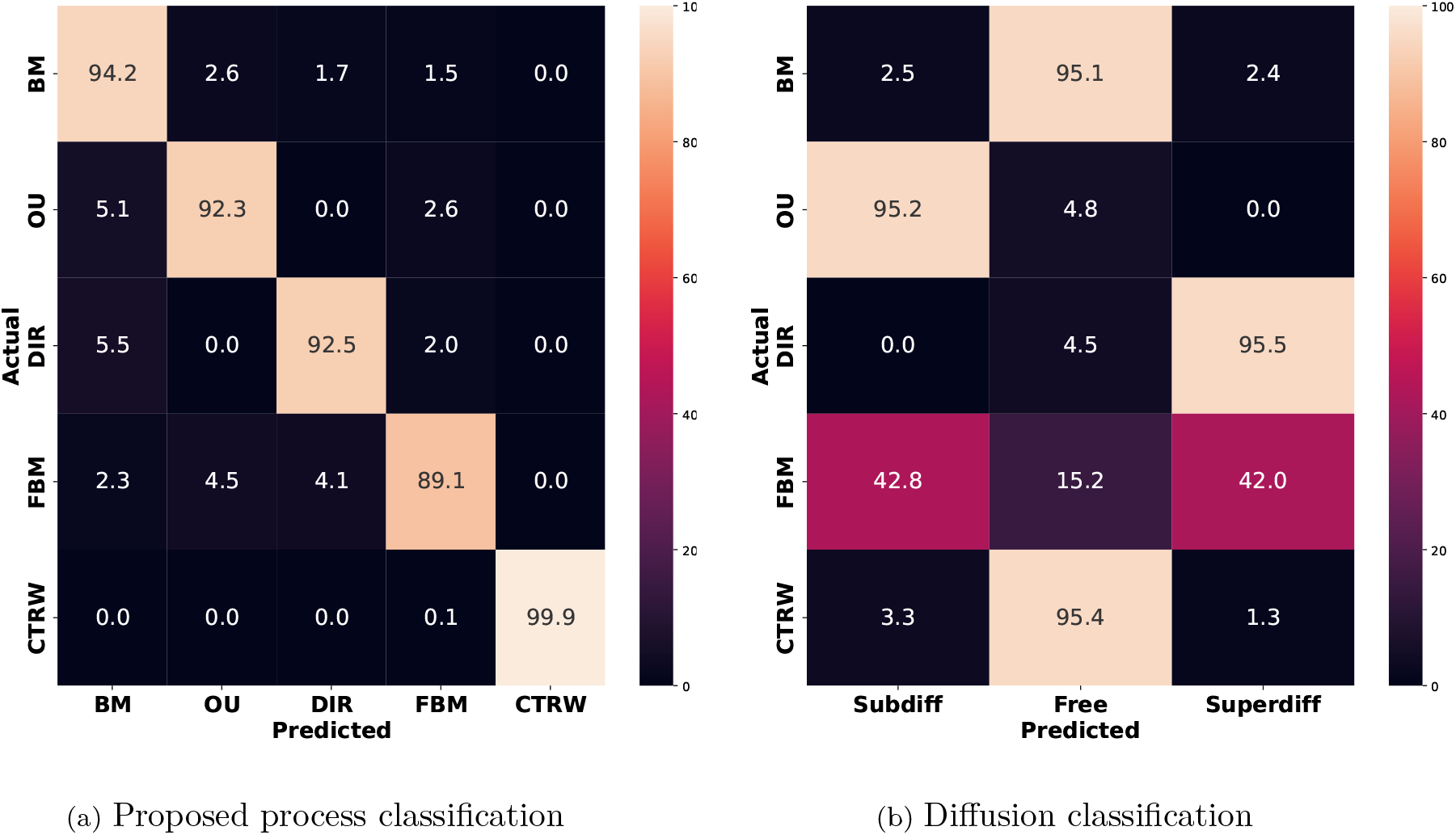
Performance comparison of machine learning and three-hypothesis-test (3.2) [9] tested on trajectories of length 100. (a) Normalized confusion matrix for the proposed model trained on trajectories of length 100. (b) Normalized confusion matrix for the diffusion classification (3.2) compared to motion labels.

Previous results show that the proposed approach, based on process classification, allows a finer description of particle dynamics, compared to the diffusion-based framework. Firstly, this enables distinguishing different subdiffusive behaviors corresponding to different environmental constraints. Secondly, this makes possible the distinction of the constrained evolution of FBM trajectories compared to OU or DIR trajectories. Finally, a precise highlighting of CTRW trajectories is guaranteed. Thus, this approach introduces a new prism for motion analysis which turns out to be extremely powerful for a finer investigation of experimental data.

#### Accounting for motion change along single trajectory

The last critical point to be addressed in tracking analysis concerns the dynamic changes in trajectories due to particle interactions with the cellular environment. It is known, for instance, that many viruses and other cargos alternate between Brownian motion and directed transport during intracellular trafficking [2, 38]. Such changes of dynamics *on the fly* are difficult to detect [39] and can lead to erroneous classification of object’s motion. To test the robustness of our method to motion change, we measure classification performance on trajectories composed of two different types of motion.

We construct composed trajectories considering the mono-dynamics trajectories from the test dataset with length *N* = 100 and composing them according to several percentages. Let *m*_1_, *m*_2_ be two types of motions, and let 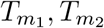 the sets of corresponding paths (1000 for each motion) in the test dataset. For a given percentage *p* and for *i* = 1, …, 1000, we consider 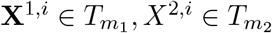 and define the following composed (bi-dynamics ) trajectory

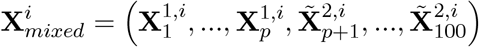

where 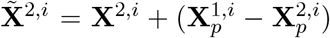. Considering several percentages (from 10% to 90%), this defines, for each *p*, a set of 1000 trajectories whose first *p* steps are governed by the *m*_1_ motion and the rest by the *m*_2_ motion. The corresponding composed trajectories are analyzed using the model trained on mono-dynamics trajectories of length *N* = 100, for each pair of composed motions and each percentage. This allows testing of how the detection of the principal motion is influenced by the type of composition and the corresponding percentage.

Figure 9 shows the main results on trajectories composed from Brownian motion and other processes, while Figure 10 concerns the other possibilities of composition. Both figures, we show confusion matrices and prediction confidence levels per composition and percentage.

**Figure 9.**
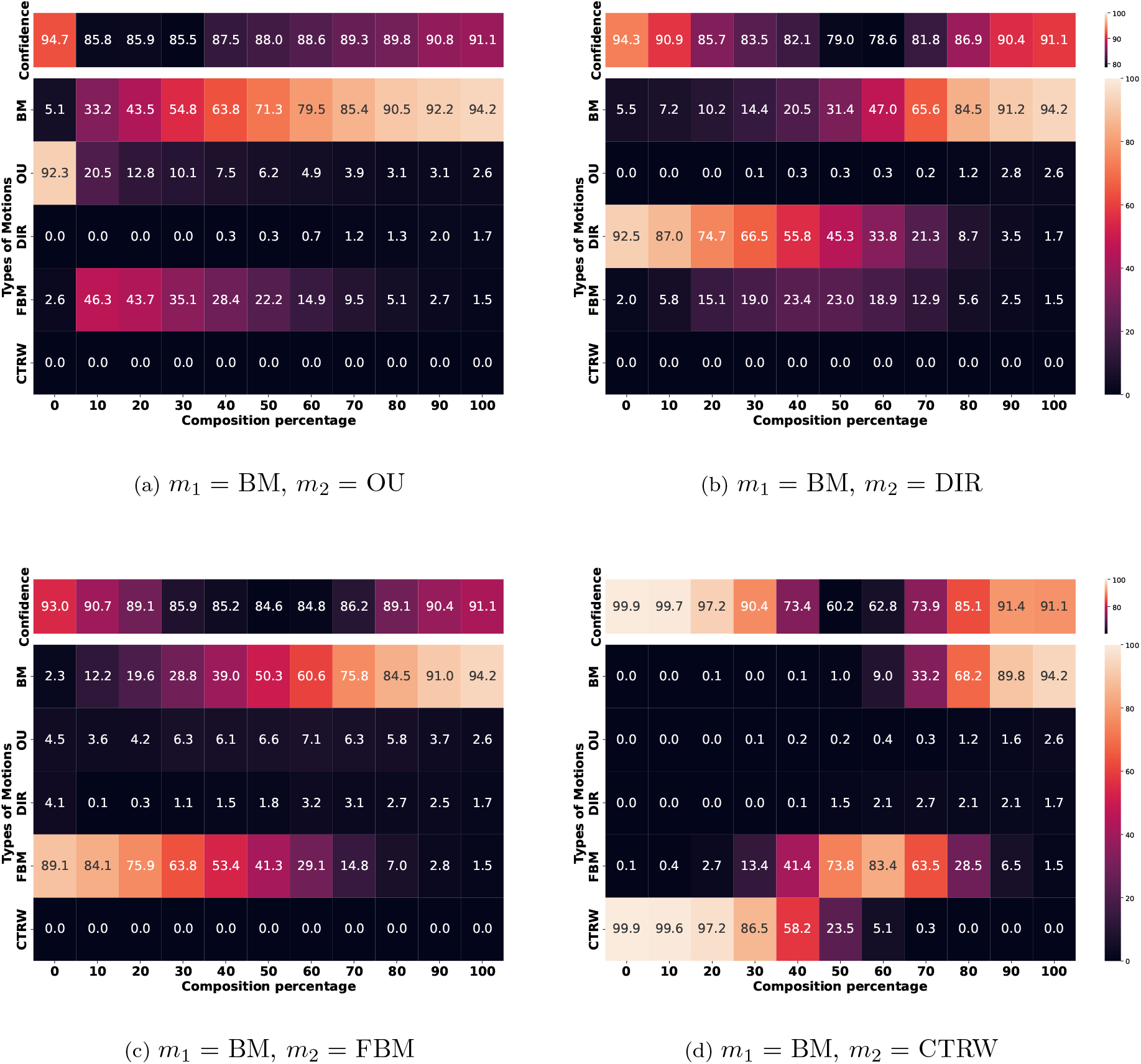
Composed trajectories involving Brownian motion with the other types of movement. The composed trajectories are analyzed with the model trained on mono-dynamics trajectories (*N* = 100, Δ*t* = 1*/*30). The recall of each class of motion and the prediction confidence level is shown depending on the composition percentage.

**Figure 10.**
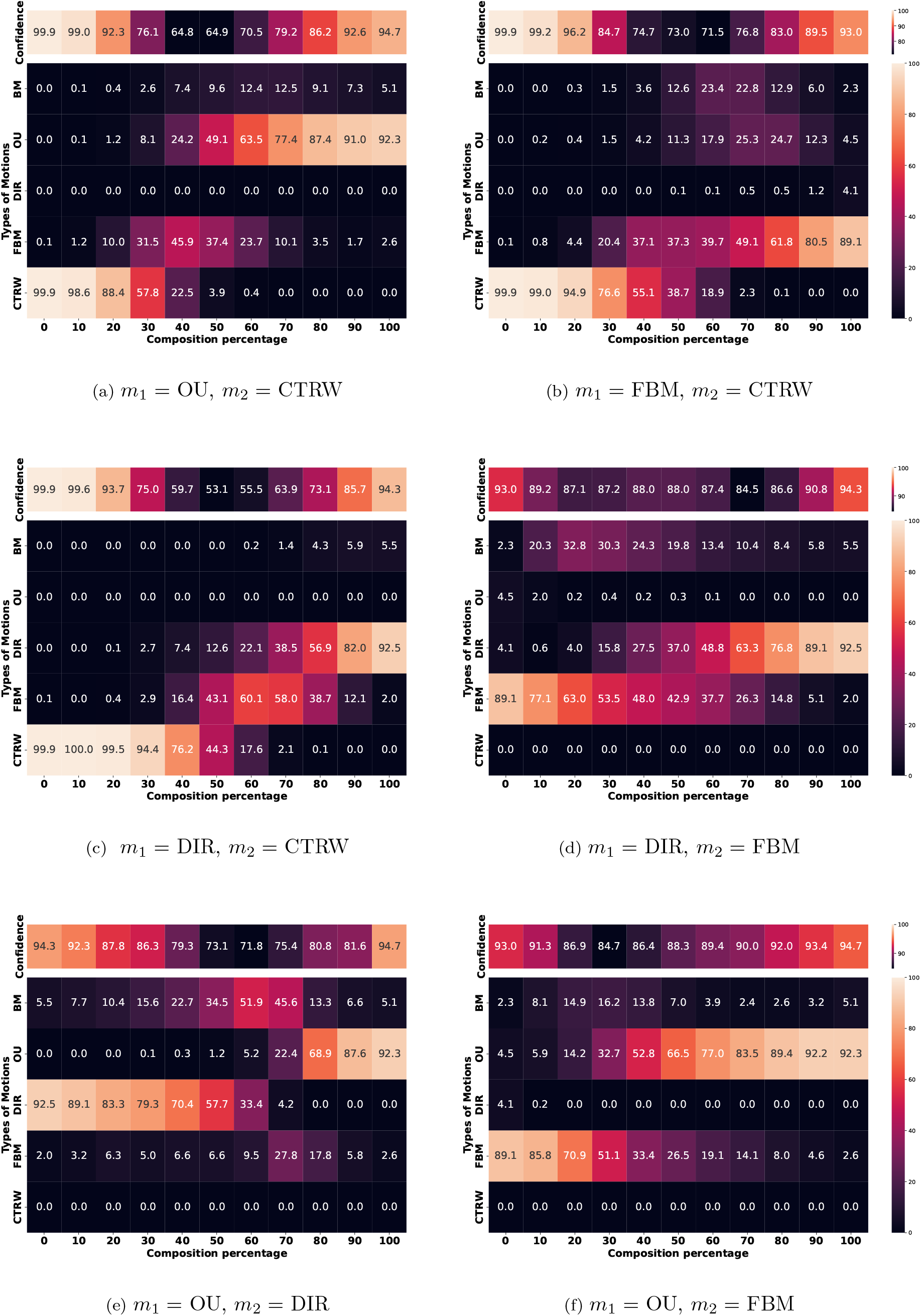
Composed trajectories involving subdiffusive and directed motions are analyzed with the model trained on mono-dynamics trajectories (*N* = 100, Δ*t* = 1*/*30). The recall of each class of motion and the prediction confidence level is shown depending on the composition percentage.

We begin by discussing the confusion matrices, displaying the recall of each class of motion as a function of the composition percentage. We point out that the method performs well for all compositions, showing a recall larger than 80% in favor of the principal motion until a percentage of 20 %. This confirms that the selected features accurately describe the considered motions and resist a non-negligible motion switching.

This is particularly true in the following cases: DIRvsFBM (Fig. 10d), FBMvsCTRW (Fig. 10b), OUvsFBM (Fig. 10f), BMvsDIR (Fig. 9b), and BMvsFBM (Fig. 9c). In fact, in these cases, the recall results show that the confusion is made essentially between the composed models, increasing for values of *p* closer to 50. This shows that the considered features are well distinguished for the related motions and confusion is due essentially to the composition percentage. This fact is particularly relevant for the case BMvsFBM showing that FBM has an intrinsic dynamics that is well distinguished from Brownian motion. Moreover, as their principal difference consists of the increments dependence, this result suggests that the selected features encode this property correctly.

A higher ratio of misclassification appears in the cases OUvsCTRW (Fig. 10a), DIRvsCTRW (Fig. 10c), BMvsOU (Fig. 9a), and BMvsCTRW (Fig. 9d). The misclassification ratio increases for the range of *p* between 30 and 70 depending on the composed motions, and interestingly misclassification always involves FBM. This seems natural for the case BMvsOU, because the OU motion introduces a more confined trend in the BM trajectory which can be globally accounted as a constrained movement with dependent increments. For the other cases, this is linked to the coupling with CTRW. We can observe that the introduction of waiting times within BM or DIR trajectories affects the particle spreading in space and the related features based on Ripley’s indices, defining global features that are more FBM-like. This fact holds also for the case OUvsCTRW for which the introduction of Gaussian-distributed jumps introduces more diffusivity in the features based on Ripley’s indices.

Finally, the most difficult case is OUvsDIR (Fig. 10e) where, for a composition percentage close to 50, the coupling of confined and directed trajectories reveals a BM behavior. This can be explained by the collection of features accounting first for a confined diffusion with large displacement angles, and then for a diffusive spreading in space characterized by small angles between successive points. The collection of well-distinguished features, corresponding to opposite motions, can explain a prediction as BM for the composed trajectory.

We now move to the analysis of the confidence tables in Fig. 9 and Fig. 10, displaying the mean confidence level for predictions as a function of the composition percentage.

We recall that Random Forest classifiers average predictions from a set of trees (ten in our case), enabling the definition of a confidence level, for the prediction of a given motion class, via the percentage of trees predicting the related process. We observe that the average confidence exceeds 90 % for almost all compositions with composition percentages up to 20 %. Moreover, this value is highly correlated with a large recall in favor of the principal motion. In particular, all mono-dynamics trajectories, corresponding to *p* = 0 and *p* = 100 in Tables of Fig. 9 and Fig. 10, present a mean confidence level larger than 91%. On the other hand, the detection of CTRW as the principal motion in composed paths shows high (*>* 91%) confidence levels until *p* = 30 (see Fig. 9d and Fig. 10a-10c).

In conclusion, previous results suggest that a low (*<* 70%) confidence level is sufficient to state the composed character of the trajectory with a percentage between 40 and 60. Although no hypothesis can be made about the nature of the composition, this adds a new powerful dimension to the method, facilitating the investigation of the link between motion switching and the biophysical constraints.

### 3.2 Characterizing CCR5 dynamics at the cell membrane

C-C chemokine receptor type 5 (CCR5) are membrane receptors that play a significant role in HIV infection. CCR5 can adopt several structural configurations, in particular different dimeric states, that interact differently with ligands or HIV envelopes and define subpopulations with distinct abilities to serve as therapeutic targets [4]. The study of their dynamic, under different conditions, has been recently initiated [4, 40] to investigate the link between motion and biological function within the infection pathways. As the studies presented in these works are based on diffusion classification the same experimental setting lends to a dynamic description in terms of stochastic process, revealing the potential of our method.

To this goal, we performed time-lapse imaging of fluorescent CCR5 in various experimental conditions. Images are acquired using TIRF microscopy, enabling visualization of a thin layer of the cell membrane (200*nm*). We collected videos at 30 fps (which is consistent with the developed method with Δ*t* = 1*/*30) by imaging several entire cells. The images are processed using the *Spot Detector* and *Spot Tracking* plugins of the Icy software [41], which allow the detection of the receptors and the reconstruction of related trajectories. Figure 11 shows the different steps of the tracking analysis on Icy. We detect spots with a size of 3 pixels (300 *nm*), and an immobility threshold of 1 pixel is applied to each path to correct positions and highlight immobility.

**Figure 11.**
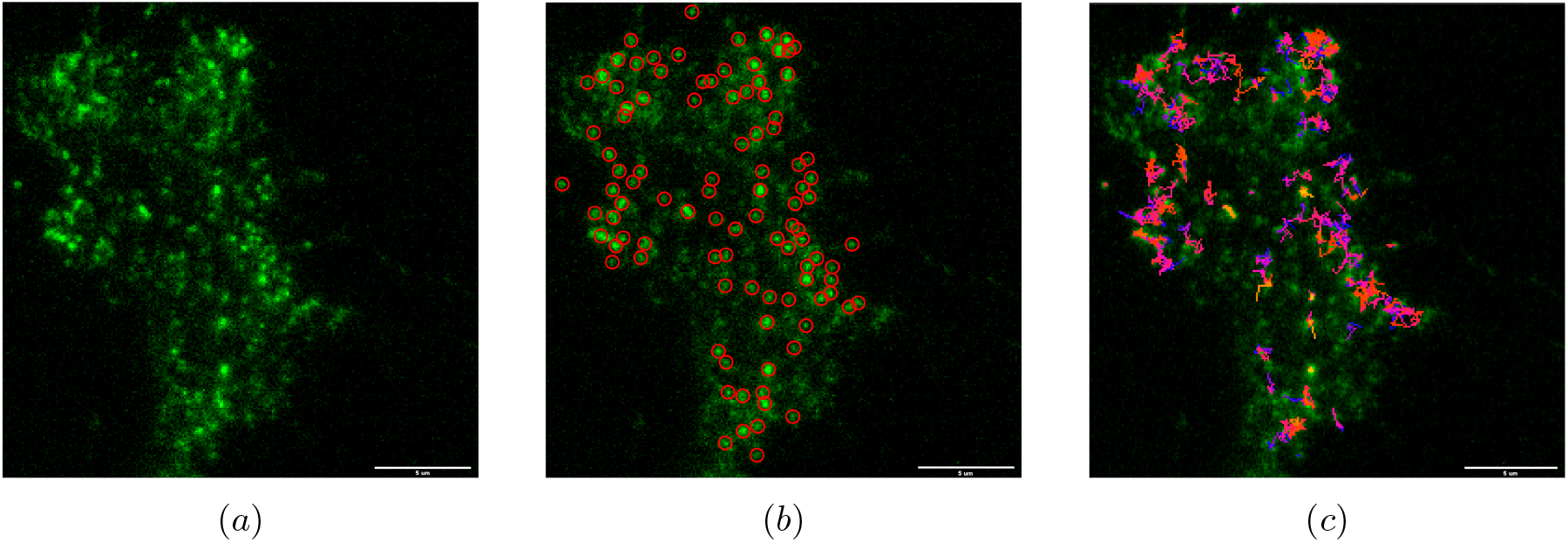
Different steps of tracking algorithm: (*a*) original image, (*b*) spot detection, (*c*) trajectory reconstruction (scale bar = 5 *µ*m).

To track CCR5 on long trajectories without ambiguities, we worked with cells expressing a low receptor level at the cell surface. We used cells expressing CCR5 under the control of the RUSH system (Retention Using Selective Hooks) developed in [42, 43]. It allows the synchronization and the study of proteins, which follow the biosynthesis/secretion pathway. It is based on the intracellular retention of a protein of interest (here CCR5) and its release by induction (using biotin). RUSH-CCR5 expressing cells, even if not biotin-induced, express CCR5 at a very low level, which corresponds to a leak from the system. The RUSH-CCR5 construct allows the cell surface expression of a CCR5 protein fused to the fluorescent protein GFP. Then, the presence of GFP-CCR5 at the cell surface is detected by labeling the cells with an anti-GFP-Atto647N booster (Chromotech). This labeling overcomes the background noise linked to the presence of GFP-CCR5 along the secretion pathways.

This experimental technique allows imaging receptors in a low-density environment, facilitating detection and trajectory reconstruction. This enables both the minimization of tracking errors and the collection of longer trajectories. We point out that we need to work with trajectories of length *N* = 100, to make the proposed method perform with high accuracy, as demonstrated in Section 2. According to the analysis of the impact of length inequality in Section 3.1, the trained model for *N* = 100 can be applied to trajectories containing between 90 and 120 time points, resulting in a final dataset of a thousand trajectories.

In the following, we perform motion classification to compare the basal state to the treatment by PSC-RANTES, an analog of the natural chemokine RANTES/CCL5, which displays potent anti-HIV-1 activity. The exceptional capacity of PSC-RANTES to inhibit infection is related to its ability to increase CCR5 down-regulation. PSC-RANTES acts as a superagonist by recognizing a larger array of CCR5 conformational states than native chemokines [44].

The results in Figure 12 reveal several motion subpopulations governed by different processes highlighting that several groups of receptors coexist, facing different environmental constraints and fates. Moreover, these results show the strong impact of the PSC-RANTES stimulation on the nature of the receptors’ dynamic.

**Figure 12.**
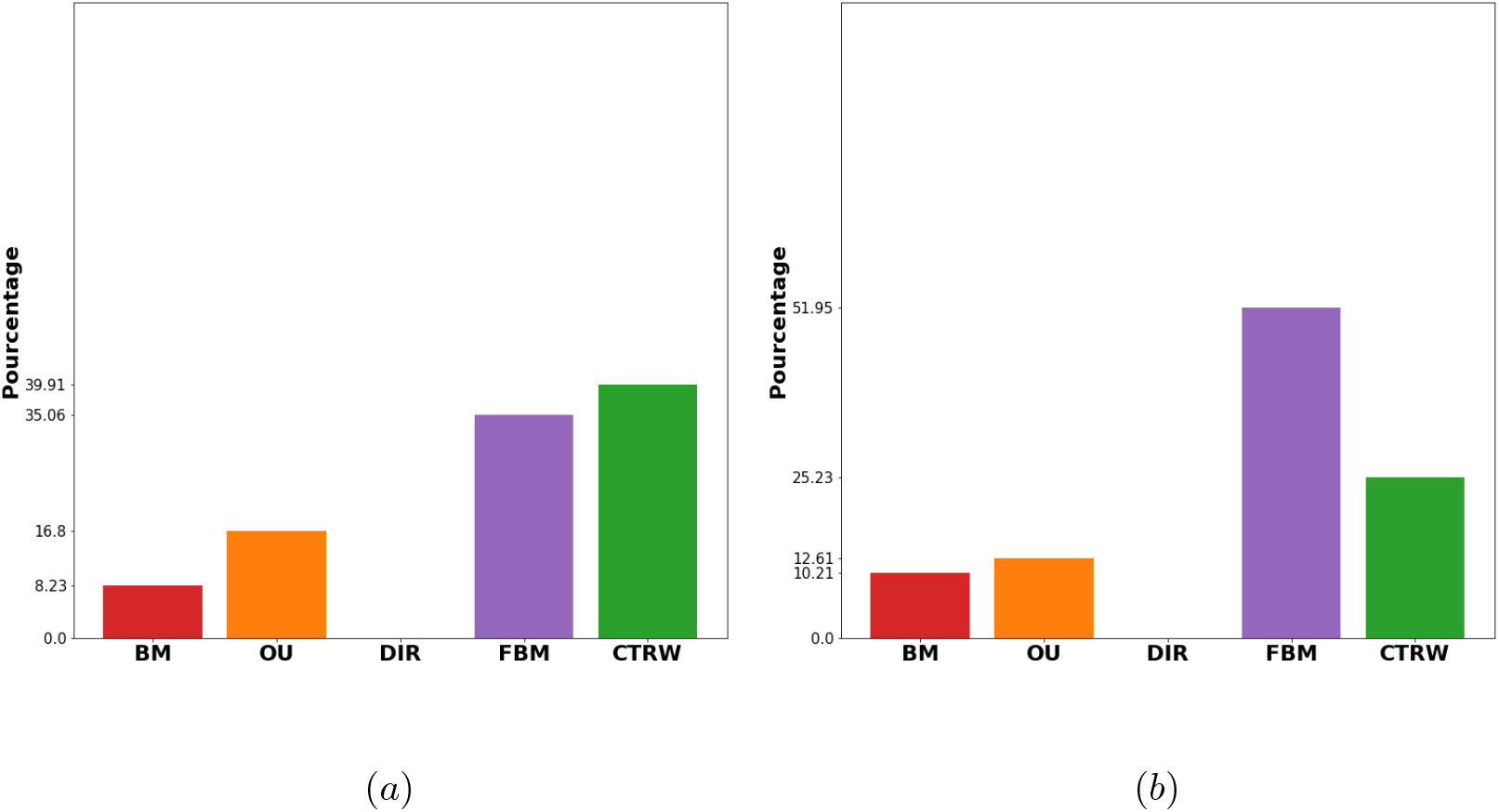
Results of motion classification for CCR5 trajectories of length approximatively equal to 100 time steps (Δ*t* = 1*/*30). Comparison between the basal state (*a*) and after PSC-RANTES stimulation (*b*).

**Figure 13.**
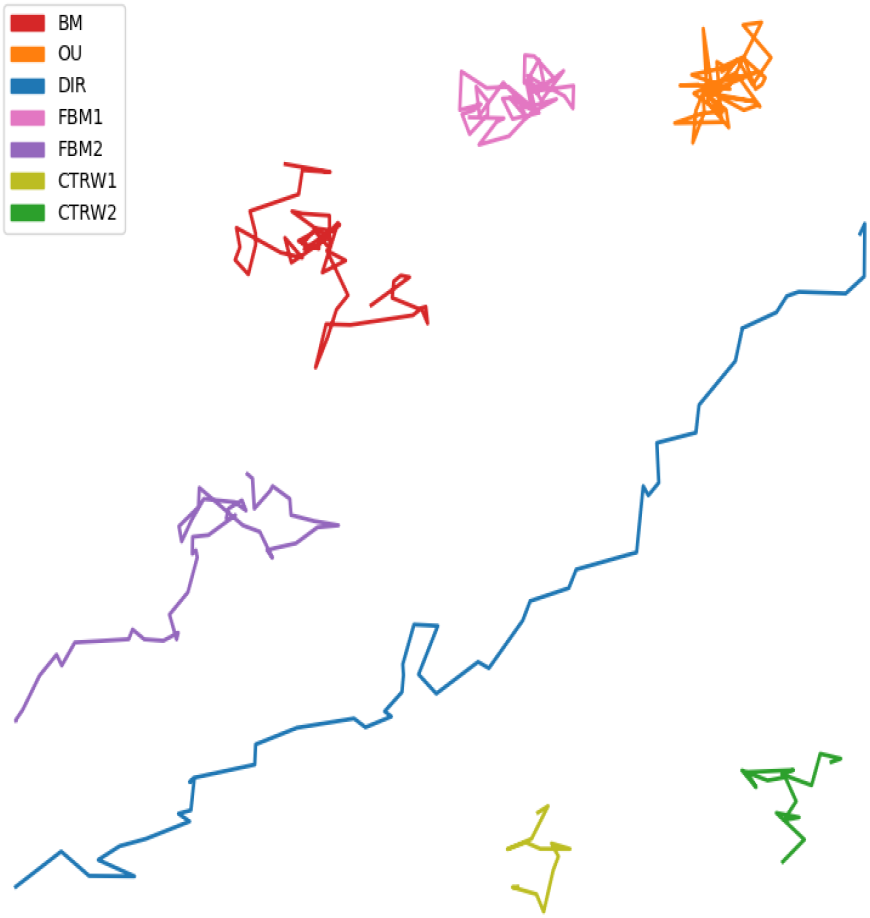
Examples of a trajectory per type of motions (*N* = 50) : BM (*σ* = 1), OU (*λ* = 0.5, *σ* = 1), DIR ( ∥*u*∥ = 0.7, *σ* = 1), FBM (FBM1 with *H* = 0.2, and FBM2 with *H* = 0.8), CTRW (*σ* = 1, *γ* = 0.01 for CTRW1 and *γ* = 0.9 for CTRW2).

This analysis defines a new powerful framework for motion classification compared to recent studies on the same topic [4, 40]. Indeed, in these works, the CCR5 dynamic is studied in terms of diffusion, highlighting a Brownian dynamic for the receptors at the basal state turning into a subdiffusive behavior after PSC-RANTES stimulation.

However, the analysis carried out in the present work reveals a more complex landscape. First of all, at the basal state, the majority of tracks exhibit either intermittent motion (CTRW) or dynamics with constrained increments (FBM). This suggests, in particular, that the Brownian motion is a too generic model to describe the natural dynamic of CCR5. Moreover, the effect of PSC-RANTES can be analyzed not only in terms of diffusion but also in terms of geometrical properties of paths. It can be seen that, after stimulation by PSC-RANTES, CTRW dynamics are less represented in favor of FBM motion. This highlights the impact of PSC-RANTES stimulation on receptors’ behavior, prompting them to move from free movement, characterized by jumps and pauses, to constrained motion at the cell membrane aiming for internalization.

In conclusion, the proposed method turns out to be a powerful lens to finely capture the CCR5 dynamics, enabling the classification of different behaviors described by distinct stochastic laws and correlating with specific environmental constraints. Thus, this allows a deeper investigation of the dynamic life of receptors, opening the door to a more precise understanding of their functional activity.

## 4 Discussion

A novel method to infer the stochastic laws (BM, OU, DIR, FBM, CTRW) governing particle dynamics has been presented. Designed to distinguish different subdiffusive behaviors, it is well suited to capture biophysical constraints in cellular environments (e.g., trapping, confinement, or constrained evolutions). The provided classification consistently refines the diffusive classes and accurately highlights the non-ergodic behavior described by CTRW. Moreover, the interpretability and explicability of predictions are guaranteed by using geometric descriptors of paths in a supervised framework. This is an additional asset of the method to emphasize the link between particle fate and dynamics geometric properties and reveal the biophysical impact of specific molecules on cell signaling. Thus, the proposed method paves a new avenue for cellular dynamics classification, and future works will be dedicated to expanding it by integrating additional stochastic laws and related geometric properties.

In addition to its high performance on simulated trajectories, the method robustly tackles the main problems associated with experimental acquisitions. Classification accuracy remains stable whenever trajectories are affected by average levels of localization error, proving that the selected features represent an intrinsic signature for stochastic processes. The method is also robust to trajectory length inequality, guaranteeing accurate classification for slightly shorter paths (up to Δ*N* = 10). These results ensure the reliability of the method’s implementation in experimental contexts. Indeed, they guarantee consistent analysis of moderately perturbed trajectories and facilitate data workflow, enabling the collection of trajectories of slightly different lengths.

Finally, the robustness to motion composition is investigated, proving that, for percentages of composition in the range of 10-20% the method accurately detects the principal process generating the trajectory. Moreover, the prediction confidence level defines an indicator suggesting the composed (mono-vs bi-dynamics) nature of paths. This information is useful for trajectory sorting (mono-vs bi-dynamics) to investigate the link between motion switching and biophysical constraints and characterize the related subsets of trajectories accordingly. Although the analysis of composed motions is a more complex problem, involving the detection of switching times and the classification o f composed motions, this is a first s tep f orward i n t he fi eld, sh owing th e po wer of machine-learning tools for in-depth analysis of particle trajectories.

The method is applied to characterize the dynamics of cell receptors CCR5 shedding new light on their dynamical behavior. The precise classification o f d ifferent subdiffusive motions highlights the impact of an agonist (PSC-RANTES) on receptor dynamics pointing out the relationship between mobility and receptor activation. This confirms the relevance of process-based classification to highlight the functional role of different ligands and drugs which is of fundamental importance for therapeutic targeting. We trust that the association of the proposed method with recent advances in live-cell imaging [45, 46] will pave the way to decipher the interplay of dynamics and biophysical constraints in cellular signaling.

## Data Availability Statement

The code and data are available at the following repository https://github.com/MotionClassification.

## Ethical Standards

The research meets all ethical guidelines, including adherence to the legal requirements of the study country. Author Contributions. Conceptualization: G.N., M.S.S., J.-C.O.-M, T.L.; Methodology: G.N; M.S.S.; Data acquisition: A.B., M.B.; Data analysis: G.N.; Writing original draft: G.N.; Manuscript editing: G.N., A.B., J.-C.O.-M, T.L.. All authors approved the final submitted draft.

Appendix A Modeling stochastic dynamics

**General definitions**. A s tochastic p rocess i s a o ne-parameter f amily *X* _*t*_ o f ℝ^2^-valued random variables defined on a probability space (Ω, F, ℙ) :

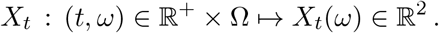

In particle tracking, the two coordinates of *X*_*t*_ are assumed to be independent random variables. The parameter *t* corresponds to the time and, for every *ω* ∈ Ω, the related trajectory is defined by the application :

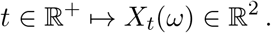

The statistical properties of increments determine the main properties of stochastic processes. A process is said to have *stationary increments* if increments distribution only depends on the time interval and not on the current times (e.g., *X*_*t*_ − *X*_*s*_ and *X*_*t*−*s*_ − *X*_0_ follow the same probability distribution for every *s < t* ∈ ℝ^+^). Similarly, a process is said to be *stationary* if the distribution of *X*_*t*_ does not depend on *t*. On the other hand, we say that a process has *independent increments* if *X*_*t*_ − *X*_*s*_ and *X*_*s*_ − *X*_*w*_ are independent random variables for every *w < s < t* ∈ ℝ^+^, implying that each displacement is not correlated to previous realizations. In the following, we consider Gaussian processes, meaning the increments follow a Gaussian distribution. Finally, we recall that a process is said to be *continuous* if it has continuous trajectories *t ↦ X*_*t*_ with probability one on Ω :

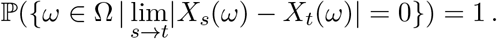

The discrete version of a stochastic process can be obtained by sampling the continuous paths on a finite s et o f t imes *t*_0_, …, *t*_*N*_ ∈ ℝ ^+^, d efining a *di screte-time st ochastic process*. A discrete trajectory is defined as a set of successive positions over time

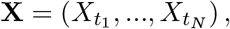

where 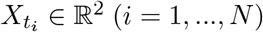 with independent components and the time interval between successive positions is constant.

### Mean square displacement criterion

The mean squared displacement (MSD) function estimates the relationship between the increments average at fixed t ime lag a nd the related time interval. Since Einstein’s seminal work on Brownian motion [11], the MSD has been used to distinguish Brownian dynamics from subdiffusive a nd superdiffusive ones [23, 24], enabling trajectory classification into three classes based on particle diffusion. However, as discussed below, this criterion is inadequate for a finer analysis aiming at distinguishing different sub diffusive dy namics or detecting non-ergodic behavior.

The MSD definition involves the ensemble average computed using the mean in the probability space [16]:

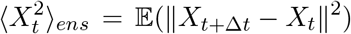

This definition can be used whenever all the particles follow the same stochastic process. Then, each trajectory represents an independent realization of it, enabling the ensemble average in the probability space, computed on the displacements of all trajectories during the time lag Δ*t*. However, in many biological contexts, particles can be governed by different processes and several dynamical behaviors can co-exist within the observed trajectories. In this case, the previous MSD definition is inadequate, a nd each trajectory needs to be analyzed individually. Then, a time-average version of the MSD function is introduced, quantifying increments average as a function of the time interval for each trajectory *X*_*t*_:

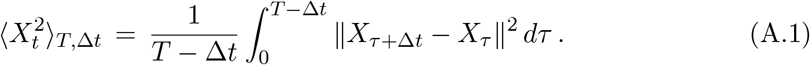

For computational applications, (A.1) can be discretized as follows :

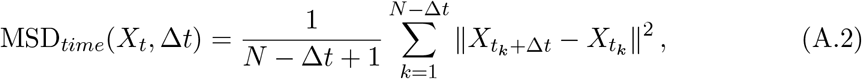

where *N* is the number of positions belonging to the trajectory **X** and Δ*t* is an integer. In single-particle analysis, trajectories are classified based on the optimal power *α* for the fitting of the time-average MSD (A.2) to the generic polynomial *t ↦ Ct*^*α*^. This enables the definition of the following MSD criterion for diffusion classification :

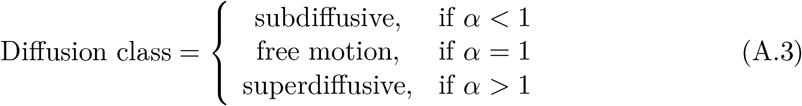

These three classes are based on particle diffusion and physically represent confined, Brownian, and spreading motions (respectively). However, this is often an inadequate and limited framework for many biological applications, for the reasons discussed below. First, a precise statistical estimate of the MSD coefficient needs long trajectories, which are often unavailable in single-particle tracking because of missed detection or temporal exit from the field of v iew. Moreover, in the case of Brownian motion, a tolerance threshold is needed to estimate the fitting coefficient *α* = 1, which introduces a new parameter to tune in the method [20].

Secondly, in the case *α <* 1, the MSD criterion detects subdiffusion, b ut i t d oes not give any information about the intrinsic behavior of the trajectory. As detailed below, subdiffusive processes can reveal different dynamics (e .g., trapping, confinement) and a more precise classification of subdiffusive behaviors is required. Moreover, beyond identifying the process governing the trajectory, estimating trajectory geometrical parameters is necessary to provide useful information on how the particle unfurls in space (for instance, privileged directions, stopping times, or spreading radius).

The last critical point about the MSD criterion concerns its consistency with ergodicity. In ergodic dynamical systems, particles visit every point of the space uniformly and randomly. This means that the statistical properties of the system can be estimated from typical trajectories rather than ensemble averages. The ergodic theorem [47, 48] states that for an ergodic process, 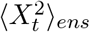and 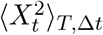 exhibit the same functional dependence on *t* and Δ, respectively. This implies that the MSD criterion based on time-average gives a result consistent with the ensemble-average estimate. However, for non-ergodic processes, time and ensemble-average MSD can exhibit different dependencies o n time, making the MSD criterion inadequate to classify trajectories governed by non-ergodic processes. This is the case, for example, of continuous-time random walks (CTRW) detailed in the next section.

### Brownian motion (BM)

Brownian motion (also called free or normal motion in some works) is a continuous Gaussian stochastic process introduced in the seminal Einstein’s work [11] on particle diffusion and widely used to model random motion. It is denoted by *B*_*t*_, with stationary and independent increments verifying :

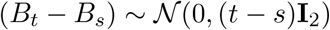

for every *t > s*, where **I**_2_ denotes the 2-dimensional identity matrix. However, this process is not stationary as *B*_*t*_ follows a Gaussian distribution with a variance depending on *t*. Setting *X*_*t*_ = *σB*_*t*_ we can vary the displacement amplitude by a parameter *σ*, and we get

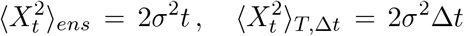

that justifies the introduction of MSD criterion. In the related literature, the MSD for BM in dimension two is written as 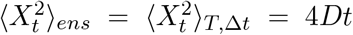 where *D* is called the diffusion c oefficient. For *X*_*t*_ = *σB*_*t*_ and, with an abuse *X*_*t*_ *σ B*_*t*_, we have the relationship *σ D* of language, *σ* is also often called diffusion coefficient.

### Superdiffusion ( DIR)

The main model for super-diffusion, which describes particles driven by active motors [49], is the so-called Directed Brownian or Directed motion and verifies

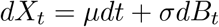

where the drift component *µ* ∈ ℝ^2^ gives a constantly oriented input to the motion. Depending on the ratio ∥*µ*∥ */σ*, the trajectory can exhibits a spreading behavior along the direction *µ* ( ∥*µ*∥ *>> σ*) or a Brownian dynamics ( ∥*µ*∥ *<< σ*). Similarly to Brownian motion, *X*_*t*_ is a non-stationary continuous Gaussian process with stationary and independent increments.

As the mean of *B*_*t*_ equals zero, a direct computation gives

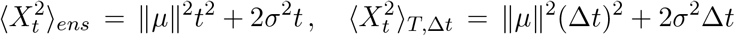

showing the superdiffusive behavior of *X*_*t*_ according to the MSD criterion.

### Ornstein-Uhlenbeck process (OU)

The continuous-time OU process [50] suits particles with limited mobility due to an external force attracting them toward an equilibrium point [32]. It verifies

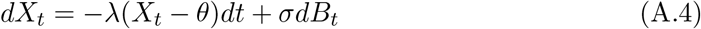

where *θ* is the equilibrium point, *σ* the diffusion coefficient of *B*_*t*_, and *λ* the weight of the drift term. In the following, up to the change of variables *X*_*t*_ = *X*_*t*_ − *θ*, we consider *θ* = (0, 0). Assuming that stochastic processes have independent components, the general results for dimension one apply.

The previous equation can be solved by variation of constants, obtaining

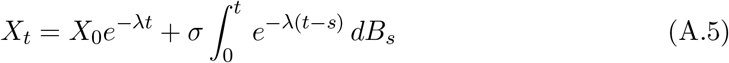

where *X*_0_ is the initial position and the integral term on right side follows the normal distribution 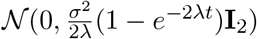. The covariance function [51] is:

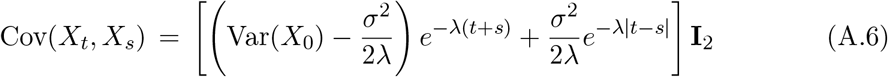

showing that the process properties strongly depend on the statistical properties of *X*_0_ (Var denotes the variance of a process).

For example, when assuming that 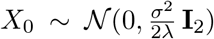 is independent of *B*_t_, we get that 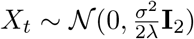, proving that *X*_*t*_ is a continuous Gaussian stationary process. The increments are also normally distributed and we have that:

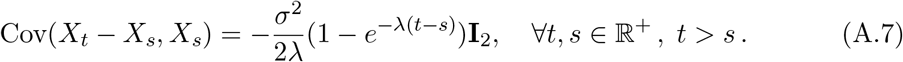

(A.7) proves that the increments are stationary but not independent because they are negatively correlated.

Otherwise, *X*_*t*_ is still a continuous Gaussian process with dependent increments, but (A.6) implies that it is not stationary (Var(*X*_*t*_) ∝ *e*^−2*λt*^) and that its increments are stationary for *t* and *s* significantly larger than 1*/λ*. This situation corresponds, for example, to paths with the same starting point *X*_0_.

In both cases, we note that the variance of *X*_*t*_ is bounded, proving that the OU process exhibits a confined diffusion: if *λ << σ*, the motion will present a Brownian-like behavior, while, if *λ >> σ*, we obtain more confined trajectories.

Finally, by a straightforward computation and using the independence of *X*_0_ and *B*_*t*_, we obtain

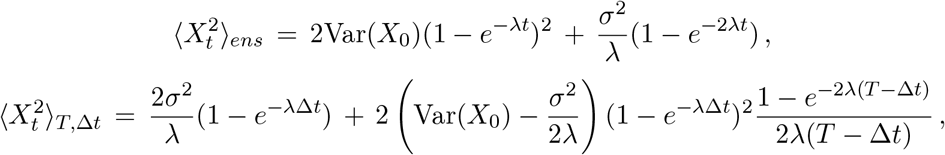

confirming t he OU p rocess’s s ubdiffusive and ergodic be havior [51].

### Fractional Brownian Motion (FBM)

It is a generalization of Brownian motion characterized by Gaussian increments that are stationary but not independent. This enables the modeling of particles moving in constrained or crowded environments [33].

Formally, a fractional Brownian motion is a continuous Gaussian process with increments verifying [52, 53, 54]:

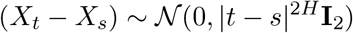

where *H* ∈ (0, 1) is the Hurst parameter which determines the intrinsic nature of this process. In particular, for *H* = 1*/*2 the fractional Brownian motion reduces to Brownian motion.

For *H* ≠ 1*/*2, as the variance of the increments depends only on the time interval and not on the current time, the process has stationary increments. However, as the variance of *X*_*t*_ − *X*_0_ depends on *t*, it is not stationary. Moreover, for *t, s* ∈ R^+^ with *t > s*, it holds

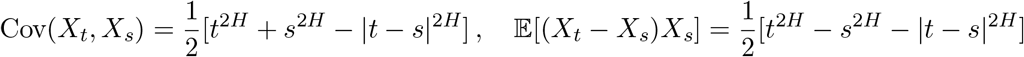

showing that the increments are not independent and that the correlation of successive increments depends on *H* (positive for *H >* 1*/*2, negative for *H <* 1*/*2).

Finally, similarly to *B*_*t*_, we obtain

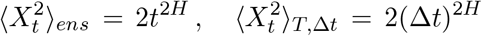

which points out the ergodicity of the process and the variety of dynamics represented by fractional Brownian motion: superdiffusive for *H >* 1 */*2, a nd s ubdiffusive fo r *H <* 1*/*2.

### Continuous-Time Random Walk (CTRW)

This process has been introduced [34] to describe trapping for random walks, exhibiting trajectories alternating jumps and waiting times; after every jump, the trajectory maintains the same position for a duration equal to the related waiting time [55, 56].

Denoting by *N* (*t*) the number of jumps up to time *t* and by the 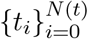 jumps times, the position of a particle at time *t* is given by:

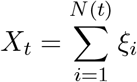

where *ξ*_*i*_ denotes the random jump and *τ*_*i*_ = *t*_*i*_ − *t*_*i* − 1_ defines the waiting time between consecutive jumps. We here assume that *t*_0_ = 0 and 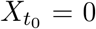. Jumps {*ξ*_*i*_}_*i*_ are independent and identically distributed (iid) random variables following a Gaussian distribution 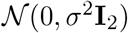, while the waiting times {*τ*_*i*_}_*i*_ are iid random variables following a power law distribution on 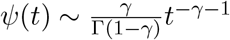 with *γ* ∈ [0, 1] and *t* ≥ 1 (see [55]). Finally, we assume that the families {*ξ*_*i*_}_*i*_ and {*τ*_*i*_}_*I*_ are independent (uncoupled CTRW).

We note that, for large *γ*, large waiting times are less likely to occur, and CTRW is then more similar to Brownian motion.

CTRW is a Gaussian process with independent increments, as *X*_*t*_ is the sum of Gaussian independent random variables. However, the variance of increments explicitly depends on the current time (see Chapter 4 in [55]), and it holds:

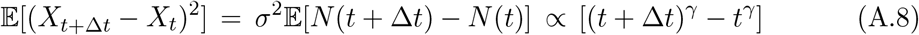

proving that the increments are non-stationary and the process is not stationary either. Furthermore, because of jumps, a CTRW path is not continuous.

Finally, CTRW exhibits the following non-ergodic property [55, 57, 58, 59]:

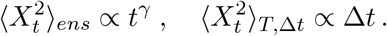

Ensemble and time averages point out different behaviors. The ensemble-average MSD is governed by the exponent of the power-law distribution for waiting times; as *γ* ∈ [0, 1], this indicates a subdiffusive t rajectory. O n t he o ther s ide, t he t ime-average M SD s uggests a normal diffusion. According to the MSD criterion, a CTRW path is classified as Brownian, pointing out that MSD method is inadequate to describe CTRW trajectories.

### Simulated trajectories

A dataset is generated for a fixed trajectory length *N*, simulating 1000 trajectories per process.

The time interval Δ*t* between two consecutive time points within the trajectory depends on the observed phenomenon and the acquisition method. The choice of this parameter has to be consistent with the acquisition process because it defines the statistical properties of several processes (for instance, the variance and covariance of increments in BM, FBM, and OU). This work considers Δ*t* = 1*/*30, a common value in confocal imaging in biology.

Brownian motion is generated by iterating random Gaussian data generation with *σ* uniformly sampled between 0.1 and 10.

An OU process is generated using *σ* between 1 and 10, and for each of them, we set 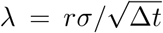 where *r* is a ratio parameter uniformly sampled between 0.2 and 1. Trajectories are simulated using the DiffusionProcess class available in the *stochastic* package of Python [60].

Similarly, directed trajectories are simulated by summing iteratively a Gaussian vector and a fixed drift vector. The diffusion coefficient *σ* is uniformly distributed between 1 and 10, the drift vector has a fixed direction (1, 1), and its norm is defined by 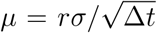 with *r* sampled between 0.2 and 1.

We point out that, considering the factor 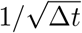 in the parameter computation for OU and DIR motions ensures that *r* represents the ratio between the standard deviations of deterministic and random increments independently of the interval Δ*t* chosen for simulations.

To simulate FBM trajectories, we use the Hosking algorithm [54], implemented in the *FBM* Python package [61], with Hurst parameters *H* uniformly sampled on [0.15, 0.3] ∪ [0.7, 0.85]. This choice enables the analysis of FBM trajectories with a behavior strongly distinct from Brownian ones (*H* = 1*/*2), which can be subdiffusive (*H <* 1*/*2) or superdiffusive (*H >* 1*/*2).

Finally, CTRW trajectories are simulated using a Gaussian distribution for jumps with *σ* uniformly sampled between 0.1 and 10. For each value, the waiting times are simulated according to an exponential distribution with parameter *γ* uniformly sampled on ]0, 1[. In particular, waiting times are sampled on the interval [1, *N/*5] to avoid too large times compared to the trajectory length *N* . Once the set of parameters is fixed, the path simulation is straightforward [56]. A sample of waiting times and jumps must be generated iteratively until the sum of waiting times is greater than *N* ; then, jumps and waiting times are alternated to define trajectories with *N* time points.

